# Single-cell sequencing reveals a clonal expansion of pro-inflammatory synovial CD8 T cells expressing tissue homing receptors in psoriatic arthritis

**DOI:** 10.1101/704494

**Authors:** Frank Penkava, Martin Del Castillo Velasco-Herrera, Matthew D Young, Nicole Yager, Alicia Lledo Lara, Charlotte Guzzo, Ash Maroof, Lira Mamanova, Suzanne Cole, Mirjana Efremova, Davide Simone, Chrysothemis C Brown, Andrew L Croxford, Sarah Teichmann, Paul Bowness, Sam Behjati, M Hussein Al-Mossawi

**Affiliations:** Nuffield Department of Orthopaedics Rheumatology and Musculoskeletal Sciences, University of Oxford, Oxford, OX3 7LD, UK; Wellcome Sanger Institute, Hinxton, CB10 1SA, UK; Wellcome Centre for Human Genetics, University of Oxford, Oxford, OX3 7BN, UK; UCB Pharma, 216 Bath road, Slough, SL1 3WE, UK; Infection, Inflammation and Rheumatology Section, UCL Great Ormond Street Institute of Child Health, London, WC1N 1EH, UK; Idorsia Pharmaceuticals Ltd, Drug Discovery Immunology, Hegenheimermattweg 91, 4123 Allschwill, Switzerland; Cambridge University Hospitals NHS Foundation Trust, Cambridge, CB2 0QQ, UK; Department of Paediatrics, University of Cambridge, Cambridge, CB2 0SP, UK

## Abstract

Psoriatic arthritis (PsA) is a debilitating immune-mediated inflammatory arthritis of unknown pathogenesis commonly affecting patients with skin psoriasis. We used three complementary single cell approaches to study leukocytes from PsA joints. Mass cytometry (CyTOF) demonstrated marked (>3 fold) expansion of memory CD8 T cells in the joints compared to matched blood. Further exploration of the memory CD8 compartment using both droplet and plate based single cell RNA sequencing of paired alpha and beta chain T cell receptor sequences identified pronounced CD8 T cell clonal expansions within the joints, strongly suggesting antigen driven expansion. These clonotypes exhibited distinct gene expression profiles including cycling, activation, tissue homing and tissue residency markers. Pseudotime analysis of these clonal CD8 populations identified trajectories in which tissue residency can represent an intermediate developmental state giving rise to activated, cycling and exhausted CD8 populations. Comparing T-cell clonality across patients further revealed specificity convergence of clones against a putative common antigen. We identify chemokine receptor *CXCR3* as upregulated in expanded synovial clones, and elevation of two CXCR3 ligands, CXCL9 and CXCL10, in PsA synovial fluid.

---

Up to one third of patients suffering from psoriasis develop the debilitating immune-mediated inflammatory joint disease, psoriatic arthritis (PsA)^1^. The pathogenesis of PsA is complex, involving multiple inflammatory pathways^2^. Genome-wide association studies of both psoriasis and PsA support a pathogenic role for CD8 T cells, showing significant associations with MHC class I and other T cell genes^3^. A subgroup of patients with PsA develop a distinct pattern of arthritis, termed large-joint oligo PsA, which affects large joints including the knees^4^. These patients often require therapeutic aspiration which provides an opportunity to examine the synovial exudate in PsA. Here, we study the cellular landscape of PsA blood and synovial fluid at single cell resolution (**Figure 1A**), combining mass cytometry with droplet-encapsulated single cell mRNA sequencing (Chromium 10x) and validation by full length transcript (Smart-seq 2) single cell mRNA sequencing (**Figure 1B**).

**Figure 1.**
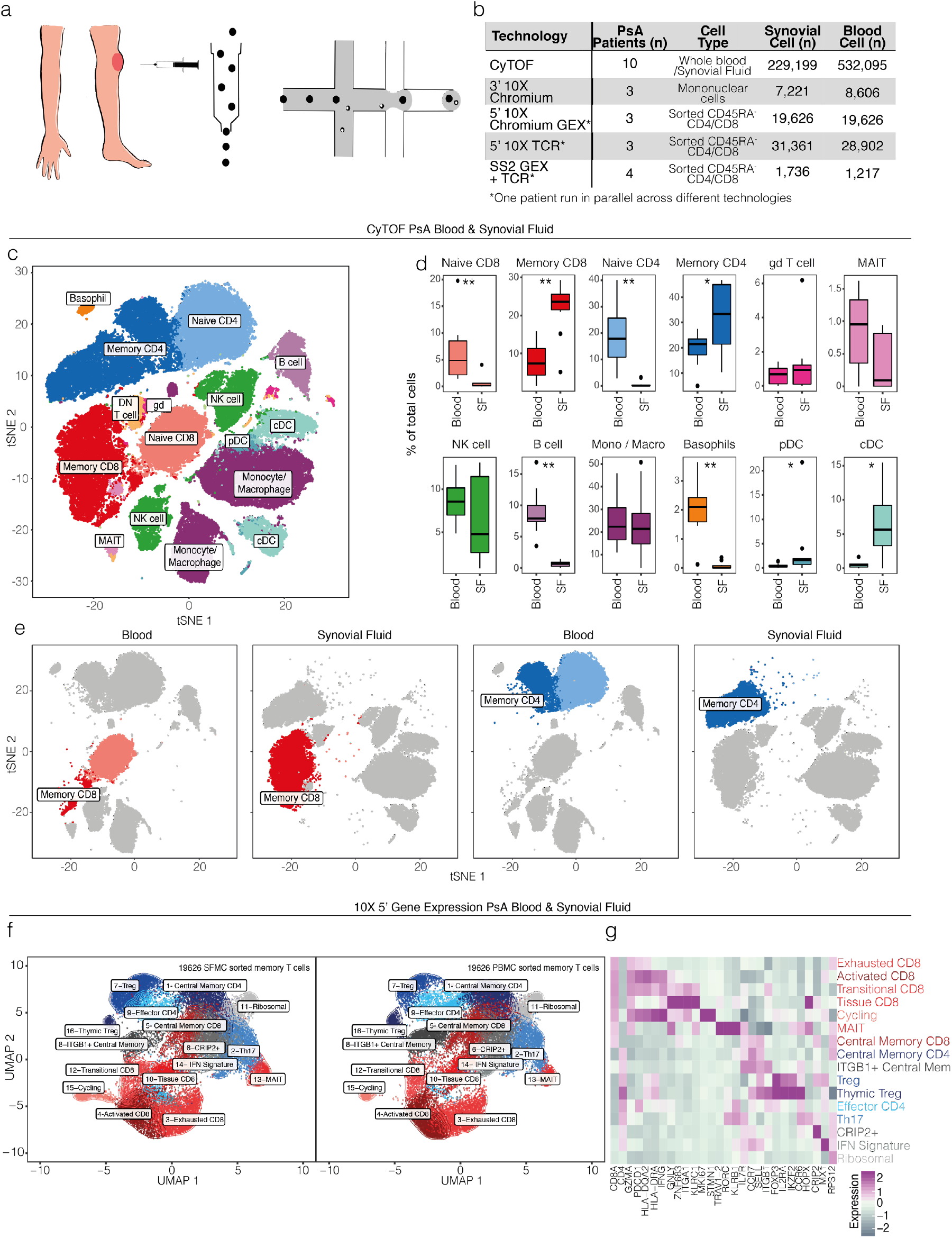
Landscape of synovial leukocyte populations in Psoriatic Arthritis. **a.** Overview of experimental design. **b.** Cell numbers used in each of the experimental techniques. **c.** Representative map of CyTOF clusters derived from PsA matched peripheral blood and synovial fluid cells using t-SNE. **d.** Cluster frequencies across blood and synovial fluid (SF), n= 10, paired t-test. (* =p<0.05, ** = p<0.01, *** = p<0.001). **e**. Representative map of CyTOF clusters divided according to tissue of origin and highlighting memory CD8 and memory CD4 T cells. **f**. UMAP of sorted synovial (left) and blood(right) memory CD4 and CD8 T cells from 3 donors after integration. Clusters coloured red are comprised of CD8 cells, clusters coloured blue are CD4 cells. Clusters in grey contain mixed CD4 and CD8 populations. **g.** Heatmap showing memory CD4 and CD8 immune subset signatures. The relative expression of marker genes (columns) across cell clusters (rows) is shown.

We first used mass cytometry (CyTOF) to quantify leucocyte populations from matched synovial fluid and blood obtained from 10 patients presenting with large-joint oligo psoriatic arthritis for arthrocentesis (**Figure 1C-D**). Blood and synovial samples were fixed within 30 minutes of collection, stained with a 38-marker panel **(Supplementary Table 1)** and then acquired on a CyTOF Helios instrument. After pre-processing, we used t-stochastic neighbour embedding (tSNE) to derive cell clusters (**Figure 1C**), which were annotated by FlowSOM^5^ (**Supplementary Figure 1)**. These analyses identified significant expansions in all patients of synovial memory CD8 (p=0.0059, paired t-test) and CD4 (p=0.025, paired t-test) T cells (**Figure 1D-E**) compared to blood. Other populations expanded in synovial fluid were plasmacytoid (p = 0.032, paired t-test) and conventional dendritic cells (p = 0.013, paired t-test) indicating increased capacity for antigen presentation. B cells and basophils were depleted (p = 0.0059 for both, paired t-test), and monocytes, gamma-delta T, mucosal invariant T (MAIT)^6^ and NK cells were unchanged (**Figure 1D**). 3’ droplet-encapsulated single cell mRNA sequencing of PBMC and SFMC from three PsA patients, carried out in parallel, confirmed the presence of these cell types and did not identify any additional cellular populations (**Supplementary Figure 1).**

To further investigate the significantly expanded memory CD4 and CD8 T cell populations (and supported by the genetic association of PsA with polymorphisms in T cell-related genes^7^), we analyzed transcription and VDJ clonality of matched synovial fluid and blood at single cell resolution. For three patients, we used droplet encapsulated single cell 5’ mRNA sequencing (Chromium 10x), with Smart-seq2 validation in four patients (applying both 10x and Smart-seq 2 technology in parallel on the same sorted cells for one donor). For both approaches, synovial fluid and blood were processed in parallel directly *ex vivo* within four hours, with single cell solutions enriched for CD4 and CD8 T cells by flow cytometry activated cell sorting (**Supplementary Figure 2**). After applying stringent quality control criteria (**Methods**), we studied 39,252 single cell transcriptomes of equal patient and tissue origin using 10x (**Supplementary table 1**), which were integrated using the Seurat 3 pipeline to derive cell clusters (**Supplementary Figure 2)**. We found 16 clusters of memory CD4 and CD8 T cells in synovial fluid and blood (**Figure 1F)**, annotated with key marker genes in **Figure 1G (Supplementary Table 1, Supplementary Figure 2).** Of note one cluster (cluster 15) derived from all three patients and predominantly composed of synovial CD8 T cells, was distinguished by high expression of the proliferation marker *MKI67*, indicating active proliferation of CD8 T cells within inflamed joints.

We next looked for evidence of clonal expansion of CD4 and CD8 T cells in the blood and synovial fluid of each of the three donors using the 10x data set (**Figure 2A-C**). For every individual we observed between 7 and 20 CD8 clones (and 1-4 CD4 clones) significantly enriched in the synovial fluid (see **Methods**, **Figure 2C**, **Supplementary Table 2**). For patient PSA1607, parallel Smart-seq 2 data showed the same maximally enriched clone with the same magnitude of clonal expansion **(Figure 2K, Supplementary Table 2).** To determine whether clonal expansion of CD8 or CD4 T cells may be driven by common antigen(s), we used the Grouping of Lymphocyte Interactions by Paratope Hotspots (GLIPH)^8^ algorithm to assess TCR complementarity-determining region 3 (CDR3) similarity and putative shared specificity across the three patients in the 10x experiment. GLIPH analysis of 40915 synovial cells with 19582 unique CDR3 beta chain amino acid sequences revealed 143 TCR convergence groups (CRG) with shared specificity between the three patients (**Supplementary table 4)**. One CRG in particular, contained a high number of cells belonging to synovial enriched CD8 clones, including the most enriched clones from 2 patients (p = 4.2E-27 and 2.1E-112 for patients PSA1505 and PSA1607 respectively), and bearing a GLIPH identified enriched CDR3 “NQNT” motif (observed versus expected foldchange 10.869, p= 0.001) relative to the expected frequency in an unselected naive reference TCR set^8^.

**Figure 2.**
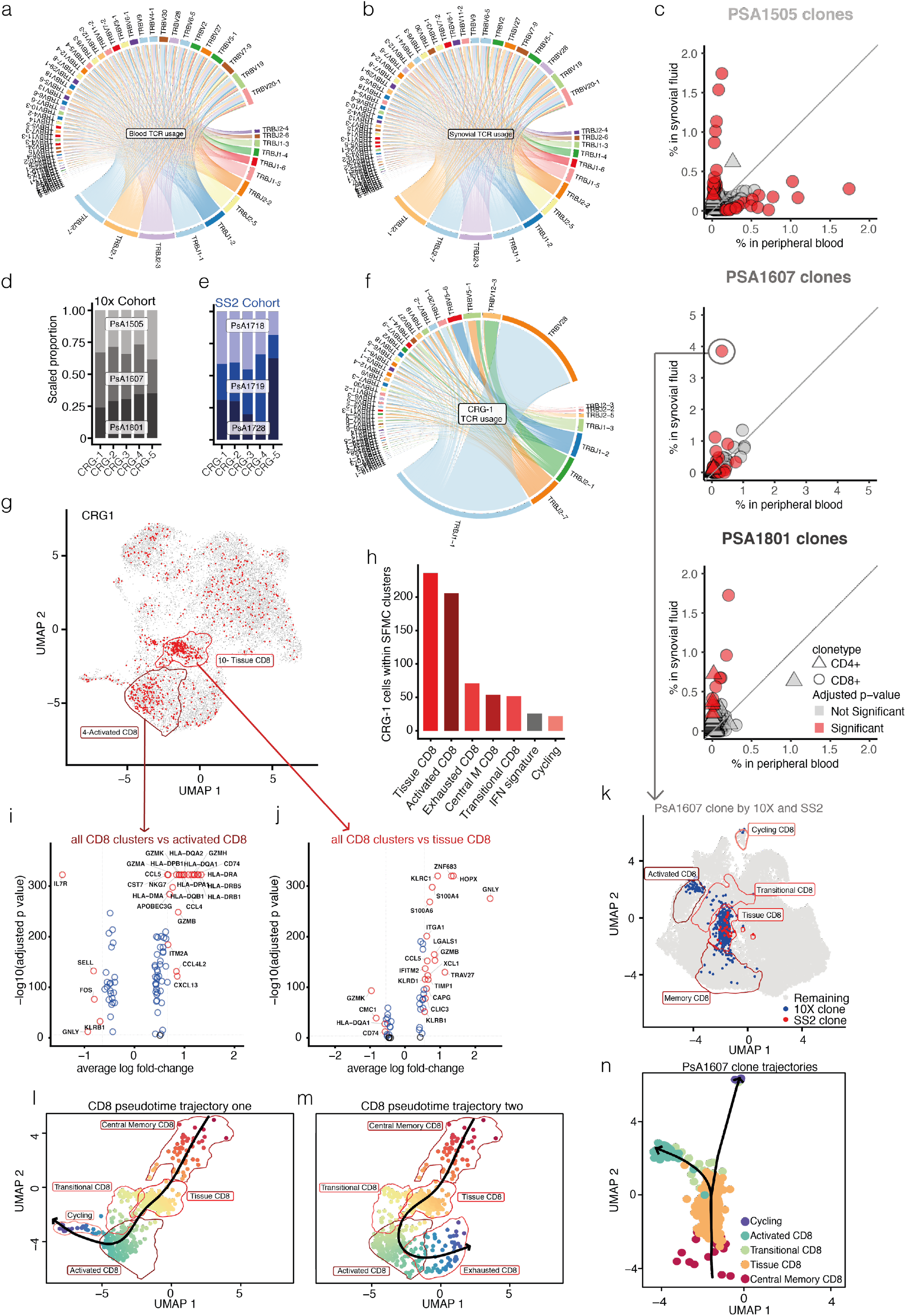
Synovial CD8 clonal expansion in Psoriatic Arthritis. **a-b.** TCR beta chain V and J gene usage in PsA blood and synovial fluid generated from 10x 5’ data. **c.** Clonal expansion across blood and synovial fluid in three PsA patients based on 10x 5’ data. Circles represent CD8 clonotypes, triangles represent CD4 clonotypes. Data points coloured red show significantly expanded clonotypes. Fisher’s exact test, Benjamini-Hochberg correction. **d.** Patient distribution of top 5 GLIPH convergence groups (CRG) in 5’ 10x data set. **e.** Patient distribution of top 5 GLIPH CRG in Smart-seq 2 (SS2) data set. **f.** TCR beta chain V and J gene usage in GLIPH CRG-1. **g.** Cells from CRG-1 highlighted on UMAP plot of all SFMC derived T cells from 5’ 10x data set. **h.** Number of CRG-1 cells within each of the synovial T cell clusters. **i.** Volcano plot showing differential gene expression of SF CD8 T cells in cluster 4 (Activated CD8) vs all other SF CD8 T cells, statistical significance calculated using Wilcoxon rank sum test. (Supplementary table 1) **j.** Volcano plot showing differential gene expression of SF CD8 T cells in cluster 10 (Tissue CD8) vs all other SF CD8 T cells, statistical significance calculated using Wilcoxon rank sum test. (Supplementary table 1) **k.** UMAP plot of integrated memory T cells from one donor (PSA1607) including cells from 5’ 10x and SS2 data sets (Supplementary Figure 3). Synovial CD8 T cells from the clone most enriched in synovial fluid for this patient is highlighted in blue for the 10x data set and red for the SS2 data set. **l.** Pseudotime analysis of CRG-1 CD8 T cells showing first trajectory differentiation pathway from central memory phenotype to tissue residency and activated states before entering cell cycle. Cells coloured by pseudotime from red to dark blue. **m.** Pseudotime analysis of CRG-1 CD8 T cells showing second trajectory differentiation pathway from central memory phenotype to tissue residency and activated states before terminating in exhausted phenotype. Cells coloured by pseudotime from red to dark blue. **n.** Pseudotime analysis of largest synovially enriched clone from donor PSA1607 showing shared central memory and tissue trajectory ending in either activated or cycling cell state (with no exhausted end state). Cells are coloured by phenotypic cluster to which they belong (Supplementary figure 3).

To validate these specificity groups we studied a further 1441 synovial CD4 and CD8 T cells with 1236 unique beta chain CDR3 amino acid sequences from three independent patients in the Smart-seq 2 data set. GLIPH analysis incorporating these sequences with the original 10x TCR sequences obtained from 40915 cells identified 5 TCR specificity groups common to all 6 patients. One of these 5 groups (CRG-1) was again assigned the “NQNT” motif and incorporated the same clones as the synovial-enriched CRG identified by droplet-encapsulated data alone (**Figure 2D-E and Supplementary Table 4**). These findings provide evidence that CD8 lymphocyte expansion in psoriatic arthritis is driven by common antigens across patients, arguing against cytokine or superantigen-driven expansion.

CRG-1 contributed the greatest number of expanded clones to the observed CD8 T cell expansions in synovial fluid and displayed high usage of the TCR genes *TRBV28* and *TRBJ1-1* **(Figure 2F)**. Of note, cells from CRG-1 were predominantly assigned to synovial cell clusters 4 and 10 **(Figure 2G-H**). Transcripts defining cluster 4 (**Figure 2I**) included granzyme *(GZMA, GZMB, GZMH* and *GZMK), CCL4, CCL5, CD74* and MHC-II, indicating an activated phenotype. Cluster 10 defining transcripts (**Figure 2J**) included *KLRC1* (NKG2A), the tissue-residency marker *ZNF683^9^*, the skin/gut homing marker *ITGA1*(CD49a) ^10^ and granulysin (*GNLY*). These two distinct CD8 clusters were also reported in a recent single-cell study of synovial tissue CD8 cells in rheumatoid arthritis^11^. When integrating 10x and smart-seq 2 gene expression data from the same patient, we observed cells from clones maximally enriched in synovial fluid from each technology within the same cluster, validating our approach **(Figure 2K, Supplementary Figure 3, Supplementary Tables 1 and 2).** The cells within CRG-1 showed a similar gene expression profile compared to their non-CRG-1 neighbours within the same cluster **(Supplementary Figure 4)**. Cells from both clusters 4 and 10 showed an overlapping gene signature with previously described tissue-resident epidermal skin CD49a+ CD8 T cells “poised for cytotoxic function” (**Supplementary Figure 4**), which were also shown to be enriched for specific TCR V gene usage including *TRBV28*^12^. Pseudotime analysis of all cells identified within CRG-1 showed two trajectories of differentiation for CD8 cells. Trajectory one (**Figure 2L**) begins with a central memory phenotype which then transitions from tissue residency to activated phenotypes before entering active cell cycle. This trajectory accounts for the clonal expansion we observe in the synovial compartment. Trajectory two (**Figure 2M**) also arises from the central memory phenotype and passes from tissue to activated states but then diverges to an exhausted phenotype. When we follow the single most expanded clone in the synovial compartment of one patient, we observe that this same clone contains cells that have predominantly central memory, tissue, transitional, activated and cycling phenotypes (**Figure 2K**). Pseudotime analysis of this single expanded clone shows the same shared central memory to transitional CD8 trajectory with either activated or cycling end states (**Figure 2N and Supplementary Figure 3)**. We propose that the CD8 tissue resident phenotype is derived from the central memory compartment and following tissue activation, can follow trajectories giving rise to activated/cycling and exhausted synovial CD8 populations. This bifurcation in T cell fate has previously been reported in response to infectious challenge.^13^

**Figure 3.**
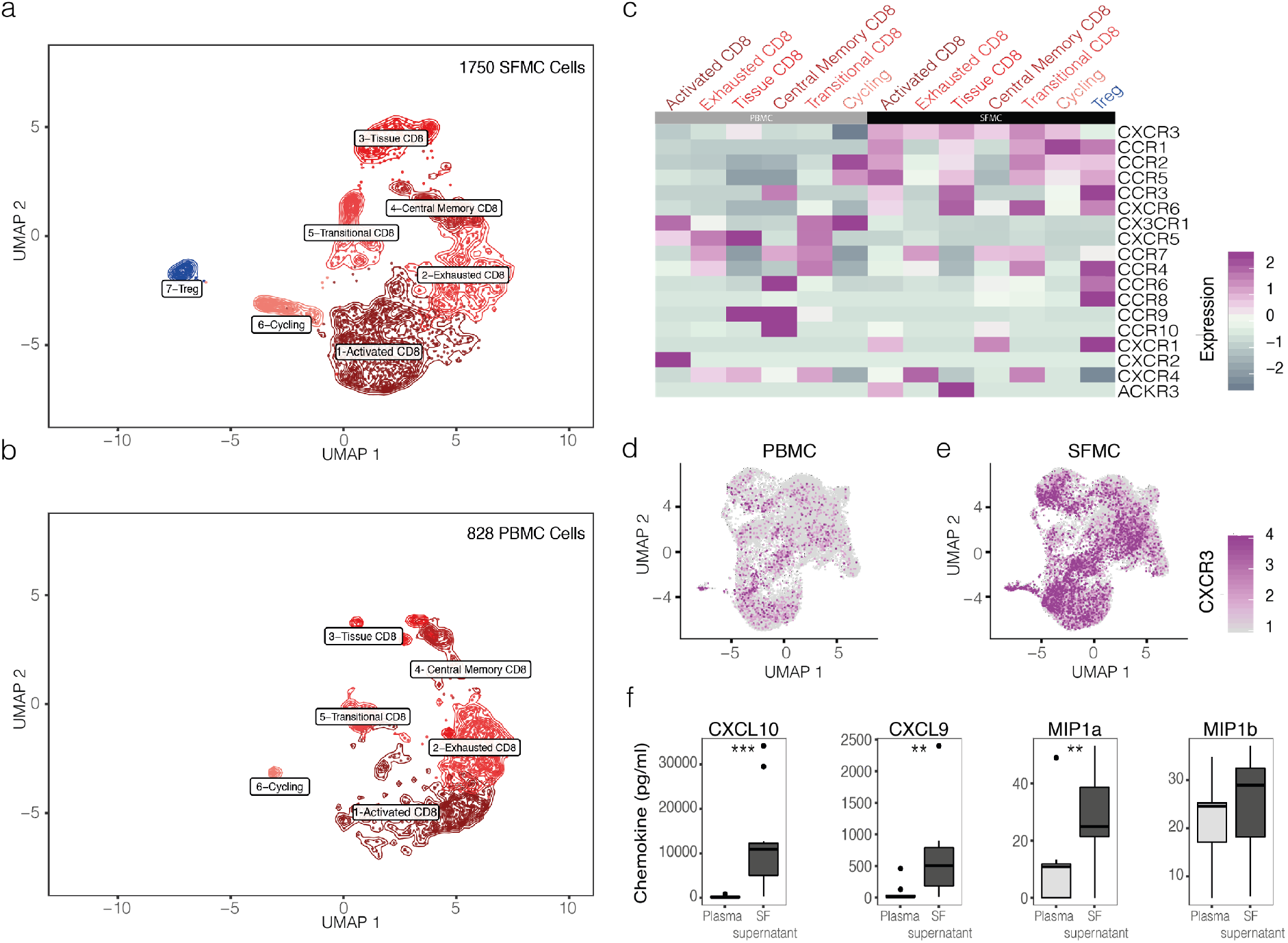
Clonal T cell trafficking in Psoriatic Arthritis. **a-b.** UMAP of 2578 (1750 synovial, 828 peripheral blood) T cells from clones significantly enriched in PsA synovial fluid and blood respectively, split by tissue of origin. Synovial cells in panel **a**, peripheral blood cells in panel **b**. **c.** Heatmap of significantly expanded blood and synovial T cell clone clusters (columns) split by tissue of origin and showing chemokine receptor expression (rows). **d.** UMAP of whole 5’ 10x data set of T cells derived from PBMC with *CXCR3* expression highlighted. **e**. UMAP of whole 5’ 10x data set of T cells derived from SFMC with *CXCR3* expression highlighted. **f.** CXCL10, CXCL9, MIP1a and MIP1b protein quantification by LegendPlex in paired plasma and synovial fluid supernatant from PsA patients, n= 10, paired t-test (** = p<0.01, *** = p<0.001).

Comparison of blood and synovial fluid TCR clonotypes showed that some clonotypes were present at both sites, highlighting inhibition of CD8 T cell trafficking as a specific therapeutic opportunity. To look for mRNAs that might mediate trafficking, we compared transcriptomes of synovial and blood T cells from clones significantly enriched in synovial fluid (n= 1,750 cells) and blood (n= 828 cells) respectively. Subsetting and re-clustering of these 2,578 cells yielded 7 clusters (**Figure 3A-C, Supplementary Table 3**) mapping back to the original clustering analysis (**Figure 1F-G)**. Genes encoding chemokine receptors *CCR1*, *CCR5*, *CXCR3* and *CXCR6* were highly expressed in synovial-enriched T cell clones (**Figure 3C**), with particularly elevated *CXCR3* gene expression (adjusted p = 7.6E-29 for activated CD8 cluster, adjusted p = 3.4E-6 for exhausted CD8 cluster, Wilcoxon rank sum test). The over-expression of *CXCR3* in synovial T cells was striking when mapped back to the whole 10x 5’ data set (**Figure 3D-E**). To functionally validate this finding, we measured protein levels of IP10 (CXCL10) and CXCL9 (ligands for CXCR3), together with MIP1a (CCL3) and MIP1b (CCL4) (ligands for CCR1 and CCR5 respectively), in the plasma and synovial fluid of patients with psoriatic arthritis (**Figure 3F**). Both CXCR3 ligands were highly enriched in the synovial fluid compared to blood (p = 0.0004 for CXCL10, p = 0.007 for CXCL9, paired t-test). CXCR3 is known to be expressed on activated Th1 and CD8 T cells and plays a key role in chemotaxis during inflammation^14,15^. Interestingly, CXCR3-expressing T cells have previously been shown to be depleted in peripheral blood of patients with psoriasis, which had been speculated to be due to recruitment of these cells into skin lesions^16^. Our findings raise the possibility that CXCR3+ CD8 T cells play a central role in executing localised inflammation in PsA and thus represent an attractive therapeutic target. Further analysis revealed that cells belonging to CRG-1 and cells from related clusters 4, 10 and 15 had selectively higher *CX3CR1* gene expression in blood **(Supplementary Figure 4**), potentially providing additional means to identify and target convergence group related CD8 T cells in blood.

In this study we have defined and characterized clonal expansions of CD8 T cells in the joints of patients with PsA, previously suggested using bulk sequencing techniques^17^, using three complementary single-cell approaches. The presence of expanded clones expressing markers of activation, tissue residency and/or tissue homing with evidence of shared TCR recognition across patients provides the strongest evidence yet that psoriatic arthritis is an MHC-I antigen driven disease. Pseudotime analysis of cells with convergent antigen specificity (CRG-1) as well as individually expanded clones provides evidence of differentiation from central memory to tissue phenotypes and insight into how this tissue-resident niche is populated in humans^18^. With application of novel approaches to identify antigens using MHC/peptide phage-display libraries^19^ or to potentially predict antigens directly from TCR sequences, these data will in future allow us to define the nature of antigens that drive psoriatic arthritis.

## Methods Summary

### Study subjects

Blood and synovial fluid samples were collected with full informed patient consent from PsA patients undergoing intra-articular aspiration at Oxford University Hospitals. The study was performed in accordance with protocols approved by the Oxford Research Ethics committee (Ethics reference number 06/Q1606/139).

### CyTOF staining and analysis

Whole blood or synovial fluid were fixed with high-purity paraformaldehyde within 30 minutes of sample collection. Fixed blood was lysed with Permeabilization Buffer (eBioscience). Cells were stained in Maxpar staining buffer (Fluidigm) with antibodies listed in Supplementary Table 1. Samples were run on a Helios system alongside normalization beads (Fluidgm). As samples were run fresh, each paired sample was analysed separately using a custom R workflow previously described^18^, with cell populations clustered using the FlowSOM algorithm^5^.

### Cell isolation for flow cytometry cell sorting

SFMCs and PBMCs were freshly isolated within 30 minutes of sample collection by density-gradient centrifugation using Histopaque (Sigma). Cells were stained immediately and FACS sorted for droplet based single cell RNA sequencing. Single cell suspensions of freshly isolated paired SFMC and PBMC samples from 3 patients were stained by a panel of fluorescently conjugated antibodies in staining buffer (RNAse-free PBS, 2mM EDTA). The following antibodies were used: CD3-FITC (SK7), CD4-APC (RPA-T4), CD8a-PE (RPA-T8), CD45RA-BV421 (HI100) (all BioLegend), together with eFluor780 viability dye (eBioscience). Memory-enriched (CD45RA negative) CD3+CD4+CD8- and CD3+CD8+CD4-cells were sorted in a 1:1 ratio from both blood and synovial fluid of patients.

### Droplet based single cell RNA sequencing

Sorted memory-enriched CD4 and CD8 T cell suspensions from peripheral blood and synovial fluid were prepared. Cells were counted and loaded into the Chromium controller (10x-Genomics) chip following the standard protocol for the Chromium single cell 5’ Kit (10x Genomics). Total time taken from sample retrieval to sample on the chip was 4 hours. A cell concentration was used to obtain an expected number of captured cells between 5000-10000 cells. All subsequent steps were done based on the standard manufacturer’s protocol. Libraries were pooled and sequenced across multiple Illumina HiSeq 4000 lanes to obtain a read depth of approximately 30,000 reads per cell for gene expression libraries and 8500 reads per cell for V(D)J enriched T cell libraries.

### Plate based single cell RNA-sequencing

Freshly sorted CD45RA negative CD3+CD4+ and CD3+CD8+ single cells from four patients were individually flow sorted into 96-well full-skirted plates (Eppendorf) containing 10μL of a 2% Dithiothreitol (DTT, 2M Sigma-Aldrich), RTL lysis buffer (Qiagen) solution. Cell lysates were sealed, mixed and spun down before storing at −80 °C. Paired-end multiplexed sequencing libraries were prepared following the Smart-seq2 (SS2) protocol^21^ using the Nextera XT DNA library prep kit (Illumina). A pool of barcoded libraries from four different plates were sequenced across two lanes on the Illumina HiSeq 2500.

### Droplet based single cell RNA-seq data mapping and pre-processing

The raw single-cell sequencing data was mapped and quantified using the 10x Genomics Inc. software package CellRanger (v2.1) against the GRCh38 reference provided by 10x with that release. Using the table of unique molecular identifiers produced by Cell Ranger, we identified droplets that contained cells using the call of functional droplets generated by Cell Ranger.

### Quality control of droplet based single cell expression data

After cell containing droplets were identified, gene expression matrices were first filtered to remove cells with > 10% mitochondrial genes, < 500 or > 3500 genes, and > 25000 UMI. Cells were further filtered to include only cells with corresponding CDR3 TCR data and to exclude potential multiplets, defined as cells with greater than 1 beta chain or 2 alpha chains or cells having both CD4 and CD8 gene expression (given the sorting strategy used).

### Quality control of SS2 based single cell expression data

Cells with more than 5 median absolute deviations (MAD) 8.35% of their mRNA originating from mitochondrial genes; a total number of reads <500,000 or > 5000000; a total number of counts >3 median absolute deviations (MAD); or with number of genes <1000 or > 6000, were filtered out prior to downstream analysis. Where matching TCR data for a cell was available, any cells with greater than 1 beta chain or 2 alpha chains were additionally filtered.

### Droplet based integrated gene expression analysis of peripheral blood and synovial fluid from 3 patients

Cellranger output included 6 expression matrices (3 patients, each with paired blood and synovial fluid samples) and downstream analyses of these matrices was carried out using R (3.6.0) and the Seurat package (v 3.0.2, satijalab.org/seurat). After quality control filtering, data were subsampled to include an equal number of cells from blood and synovial fluid from all patients (6542 cells per sample, totaling 39252 cells). All 6 data sets were then individually normalised and variable genes discovered using the sctransform function with default parameters, providing total number of UMIs and mitochondrial fraction as factors to regress out. The FindIntegrationAnchors function command was subsequently run with default parameters (dims = 1:30) to discover integration anchors across all samples. Any TCR genes were excluded from anchor features discovered prior to downstream analysis, and the IntegrateData function was run on this reduced anchorset with default additional arguments. ScaleData and RunPCA were then performed on the integrated assay to compute 30 principal components (PC). Uniform Manifold Approximation and Projection (UMAP) dimensionality reduction was carried out and Shared Nearest Neighbor (SNN) graph constructed using dimensions 1:30 as input features and default PCA reduction. Clustering was performed on the Integrated assay at a resolution of 0.7 with otherwise default parameters which yielded a total of 16 clusters, each composed of cells originating from both blood and synovial fluid from all 3 patients and classified by differentially expressed genes. (**Supplementary Figure 2).**

To compare synovial (1750) and blood (828) cells from synovial and blood enriched clones respectively, all significantly enriched clones from both tissues were isolated from the previously filtered integrated data set using the SubsetData function. The integrated assay of this subset was scaled using the ScaleData function before running principal component analysis. Dimensionality reduction was calculated with UMAP using the first 20 principal components and clustering set at a resolution of 0.4 to reflect the smaller cell numbers relative to the parent data set. This yielded 7 clusters.

### Droplet and SS2 based integrated gene expression analysis of PSA1607 peripheral blood and synovial fluid

For validation of sequencing data across platforms, quality control filtered 10x expression matrices from 1 patient (PSA1607) were subsampled to include an equal number of cells from blood and synovial fluid (11192 cells from each, totaling 22384 cells). Quality control filtered SS2 data matrices from this same patient were similarly subsampled to include 433 cells from both blood and synovial fluid. The same steps outlined above for integration of droplet based only data sets from 3 patients were used to integrate these 4 data sets (blood and synovial from both 10x and SS2 platforms) for patient PSA1607. Apart from additionally regressing out platform when scaling data, the same optional arguments specifying 30 principal components and a cluster resolution of 0.7 were used. This yielded 13 clusters which were again classified by differentially expressed genes, bearing gene expression signatures that overlapped with clusters identified in the 10x only integration of 3 patients **(Supplementary figure 3, Supplementary Table 1)**.

### TCR mapping

Chromium 10x V(D)J single-cell sequencing data was mapped and quantified using the software package CellRanger (v2.1) against the GRCh38 reference provided by 10x Genomics with that release. The generated consensus annotations files for each patient and sample type (blood or synovial fluid) were then used to construct clonality tables and input files for further downstream analysis. Full-length, paired T cell receptor (TCR) nucleotide sequences from SS2 data were constructed using the TraCeR program^22^, and further mapped to V, D and J genes as well as CDR3 nucleotide and amino acid sequences using the online IMGT-HighVQuest tool^23^. The software package GLIPH v 1.0 was used to construct and assess specificity groups^8^. As the GLIPH algorithm only makes use of single CDR3 beta chain amino acid sequences to associate clones of common specificity, multiple beta chain sequences within the same partition were treated as multiplets and provided as separate individual sequences to GLIPH (annotated with a “v” suffix). Partitions containing only alpha chain sequences were excluded from GLIPH input. The VDJtools 1.1.8^24^ software package was used to construct circle plots illustrating V(D)J gene usage.

To assess clonal enrichment, the proportion of cells having the same clone was compared between sample types for each clone using a Fisher’s exact test with Benjamini & Hochberg (1995) correction for multiple comparisons (R Stats Package). A cell’s clonotype was defined as the combined alpha and beta chain CDR3 nucleotide sequences for that cell. As it was not possible to deduce beta and alpha chain pairing for partitions with multiple beta chains, these partitions were treated as a single clone.

### Plate based single cell RNA-sequencing expression quantification

To assess the expression from the SS2 data, raw reads were pseudo-mapped and counted using kallisto v0.43.1^23^ based on the annotation made by ENSEMBL(v90) of the human reference genome (GRCh38). To obtain per gene counts, all of the transcript counts were summarised using scater v1.6.3^26^.

### Chemokine protein quantification

Paired plasma and synovial fluid supernatant was collected from patients undergoing knee aspiration procedures and frozen at −80°C within one hour of sample collections. Samples were thawed and chemokine protein quantification was performed using a LEGENDplex™ Human Proinflammatory Chemokine Panel (13-plex) immunoassay (Cat# 740003) according to the manufacturer’s instructions. Samples were acquired on two different dates, with similar results obtained. The data was acquired on a Novocyte instrument and analyzed using the LEGENDplex™ Data Analysis Software provided by BioLegend

### Statistical Analyses

Statistical tests were performed as indicated in the figure legends.

### Data availability

The raw sequencing data generated for the present study has been deposited in the European Genome-phenome Archive.

## Acknowledgements

We thank Jon Webber and Catherine Simpson for assistance with cell sorting. We are grateful to Muzlifah Haniffa for critical review of the manuscript. The research was funded by The Kennedy Trust Studentship (FP), The Academy of Medical Sciences Grant to HA (SGL018\1006) and Personal fellowship support HA (Oxford-UCB Prize fellowship) SB (Wellcome; St Baldrick’s Foundation), CB (Wellcome). PB and HA are also funded by the National Institute for Health Research (NIHR) Oxford Biomedical Research Centre (BRC). The views expressed are those of the author(s) and not necessarily those of the NHS, the NIHR or the Department of Health.

## Author Contributions

HA-M and SB conceived and designed the experiments; FP and HA-M performed the 10x experiments, designed and performed computational analysis aided by MDCVH, MDY, LM, ME and DS; MDCVH performed the SMART-seq2 experiments assisted by CG and LM; NY performed the CyTOF experiments assisted by ALL, SC and AM; NY, ALL and SC performed flow cytometry and assisted with cell-sorting; ALC performed protein quantification assays; ST and CB contributed to discussions; HA-M, SB and PB wrote the manuscript, PB, SB and HA-M co-directed this study.

## Author Information

The authors declare no conflict of interest. Correspondence and requests for materials should be addressed to.

